# Examining the early distribution of the artemisinin-resistant *Plasmodium falciparum* kelch13 R561H mutation in Rwanda

**DOI:** 10.1101/2022.10.26.513523

**Authors:** Rebecca Kirby, David Giesbrecht, Corine Karema, Oliver Watson, Savannah Lewis, Tharcisse Munyaneza, Jean de Dieu Butera, Jonathan J Juliano, Jeffrey Bailey, Jean Baptiste Mazarati

**Author notes:** Correspondence should be made to Jeffrey Bailey or Jean Baptiste Mazarati. Authors contributed equally (Rebecca Kirby, David Giesbrecht, Jonathan J Juliano, Jeffrey Bailey, Jean Baptiste Mazarati).

## Abstract

Artemisinin resistance mutations in *Plasmodium falciparum kelch13* (*Pfk13*) have begun to emerge in Africa. *Pfk13-*R561H was the first reported African mutation found in Rwanda in 2014, but limited sampling left questions about its early distribution and origin. We detected 476 parasitemias among 1873 residual blood spots from a 2014-15 Rwanda Demographic Health Survey. We sequenced 351 samples revealing 341/351 were wild type (97.03% weighted) and 4 samples (1.34% weighted) harbored R561H which were significantly spatially clustered. Our study better defines the early distribution of R561H in Rwanda and suggests that the origin may have involved higher-transmission regions.

## Introduction

Artemisinin-based combination therapies (ACT) are currently the key antimalarials used in Africa and emerging resistance threatens decades of public health gains. Concerningly, mutations in the *Plasmodium falciparum kelch13* (*Pfk13)* gene causing in vitro artemisinin resistance are emerging in Africa [1,2]. Additionally, recent reports have shown delayed parasite clearance in Rwandan patients infected with R561H mutants after treatment with ACT [3]. While high-grade clinical resistance to ACT remains low in Africa thanks to the continued efficacy of partner drugs, decreased sensitivity to artemisinin is thought to have potentiated partner drug and lead to ACT treatment failure in southeast Asia (SEA), making the emergence of validated *Pfk13* resistance mutations a grave concern [4–6].

Early in the emergence of artemisinin resistance in SEA, multiple mutations in *Pfk13* were found with the eventual emergence of C580Y as the dominant genotype. A similar pattern may be occuring in Africa with multiple validated or candidate resistance mutations emerging including R561H, C469F/Y, and A675V [1,2,7,8]. Understanding trends in resistance allele frequencies can only be done through intensive genomic surveillance.

The first report of artemisinin resistant *Pfk13* mutants in Africa occurred in Rwanda in 2014, with 7.4% of samples collected in Masaka between 2014-15 harboring R561H [1]. By 2018, reported R561H prevalence increased to 19.6% in Masaka and 22% in Rukara [9]. While ring-stage survival assays confirmed in vitro artemisinin resistance, these studies found no association between clinical outcomes and *Pfk13* mutations. The expansion of a validated resistance mutation is nevertheless concerning, and the lack of broader sampling in previous studies makes it difficult to trace R561H’s spread through the country. To our knowledge, only two *Pfk13* genotyping studies were done in Rwanda between 2010-2015, the years preceding and including the first report of R561H [1,10] (Figure 1A). These studies were limited to a few sampling sites, leaving questions about the true distribution of R561H early in its emergence.

**Figure 1:**
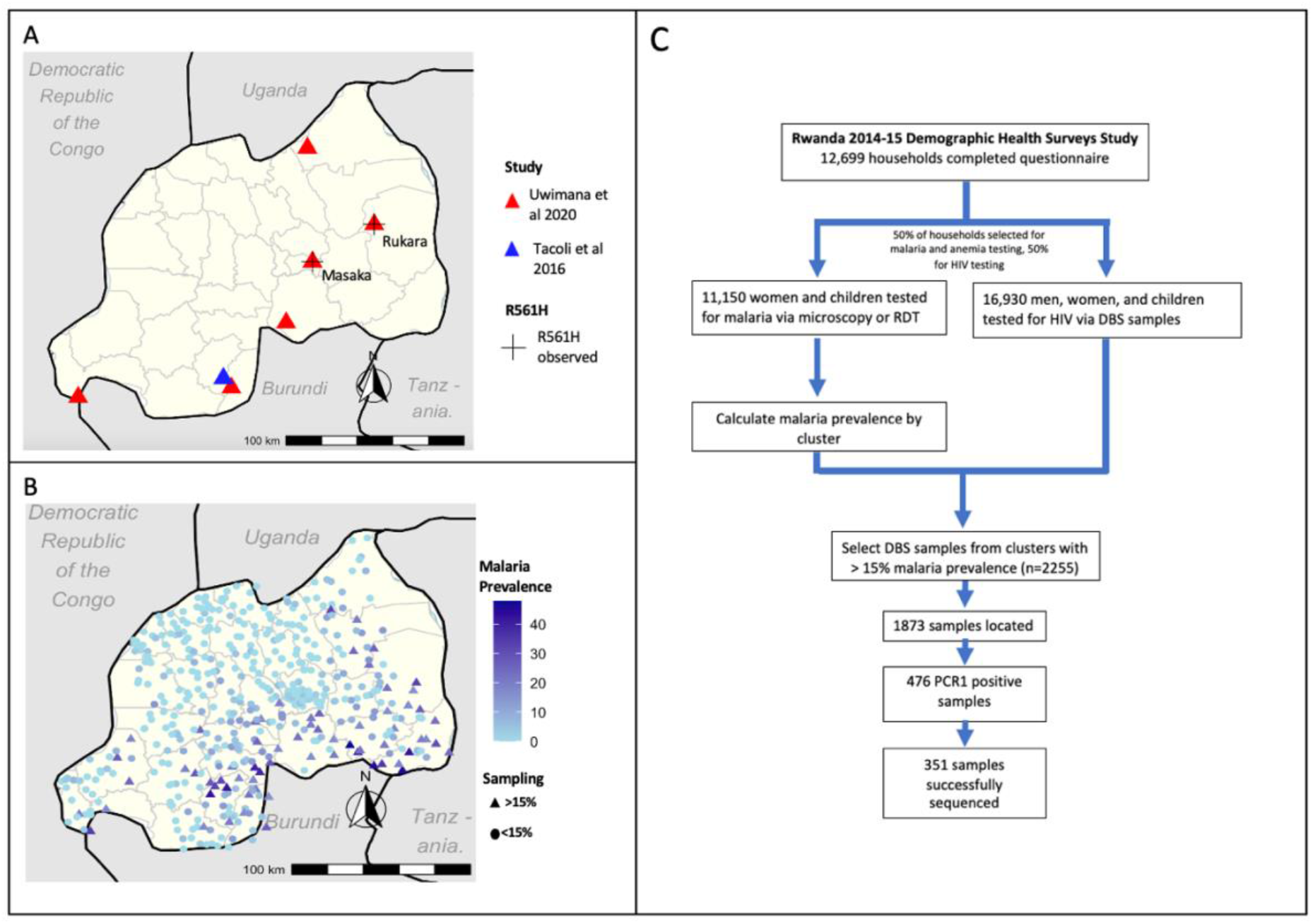
Previous Studies and Our Study Design. **A:** Sampling sites of P. falciparum genotyping studies done in Rwanda 2010-2015. Sites included Masaka, Ruhuha, Bugarama, Kibirizi, Nyarurema and Rukara (Uwimana et al 2020) and the Huye district 2010-15 (Tacoli et al 2016). **B:** DHS 2014 sampling clusters. We sequenced samples from clusters with >15% malaria prevalence. **C**: Study design. We used DBS samples from the HIV arm of a 2014-15 DHS study, selecting 2255 samples taken from 67 clusters with >15% malaria, as calculated in the malaria/anemia arm.

Here we used an existing Demographic Health Surveys (DHS) sample set, originally designed as a random representative sample of the Rwandan population for HIV prevalence, conducted between 2014-15. Repurposing this nationally representative dataset provides an opportunity for a nation-wide baseline study of R561H prevalence in Rwanda early in the emergence timeline (Figure 1B). This study also employs a cost-conscious approach for genotyping discarded samples that can be employed in resource-limited settings.

## Methods

### Sample collection and preparation

Sample collection for 2014-15 DHS in Rwanda included a subset of participants from whom dried blood spots (DBS) were collected on filter paper for HIV testing. Residual DBSs were provided by Rwanda Biomedical Center. The DHS included 12,699 GPS tagged households in 492 clusters in 30 regions who completed the initial questionnaire. Of these households, 50% were selected for malaria and anemia testing in women and children, and the other 50% were selected for HIV testing via DBS sample collection (Figure 1C). Complete survey methods are described by the DHS [11]. The final anonymized DHS dataset was downloaded from https://dhsprogram.com. All data cleaning and analysis was done in R version 4.1.2 using packages detailed here: https://github.com/bailey-lab/Rwanda-DHS-2014-15. Analyses utilizing parasite genomes from de-identified samples were deemed nonhuman subjects of research at the University of North Carolina at Chapel Hill (NC, USA) and Brown University (RI, USA).

From the DHS survey database, malaria prevalence for each cluster was calculated based on either positive rapid diagnostic test (RDT) or microscopy. Given we could not simply identify positive DBS samples based on RDT testing as they were different groups, we focused on clusters where we would expect to detect a reasonable number of positives to genotype, selecting clusters with >15% malaria prevalence (n clusters = 67, n samples = 2255). Regional malaria prevalence was calculated in R using survey-weighted means, using DHS household survey-weights. We located 1873 of these samples and placed three 6 mm punches into wells of a 2 ml plate. DNA was extracted using Chelex [13].

### Amplification of short PCR amplicons including R561H site

A two-step PCR was optimized to amplify short amplicons with sample barcodes covering amino acid positions 516-572 on the Pf*k13* gene (Supplementary Methods). PCR step 1 product was resolved on an agarose gel, and the presence of a faintly visible or stronger band was used to select samples for sequencing. First-step primers were based on previous designs [12] and contained molecular inversion probe (MIP) compatible 5’ linking sequences, enabling barcoding [13]. Positive samples (n=476) were re-plated using an OT2 robot (Opentrons, Brooklyn, NY, USA). PCR step 2 was used to add unique barcodes to each sample, and these products were resolved on a gel.

### Library preparation

To reduce over sequencing of high parasitemia samples, 5 uL of each PCR step 2 reaction were sorted into high and low parasitemia pools based on the PCR step 2 band intensity, using an Opentrons OT2. To prepare libraries for sequencing, a 1.2x SPRI bead cleanup was performed on the two pools eluted in 30uL TE low EDTA pH=8 (Thermo Fisher Scientific, Waltham MA, USA). The eluted pools were combined for a total volume of 20 uL, and amplicons were resolved on a 1.0 % agarose gel. Fragments of the correct size were excised and DNA was purified using the NEB DNA Gel Extraction kit T1020S (Ipswich, MA, USA). Library quality was assessed using a fragment analyzer (Agilent, Santa Clara, CA), the library pooled with other libraries and sequenced on an Illumina Nextseq 550 using a 300-cycle mid output kit at 5.3 pM.

### Analysis

Samples were demultiplexed using MIPtools software available at https://github.com/bailey-lab/MIPTools and analyzed using SeekDeep pipeline available at https://github.com/bailey-lab/SeekDeep [14]. Samples that had < 50 reads on the first pass of sequencing (n=243) were repooled, gel extracted, and resequenced. Samples with ≥ 20 total combined reads were included in subsequent analysis (n=351). Mutation prevalence was calculated using HIV survey-weights.

Spatial correlation of R561H was calculated in GeoDa (https://geodacenter.github.io) using Moran’s I with Empirical Bayes to take varying cluster sample sizes into account. Significance was calculated in Geoda with 999 permutations. A 97.5% confidence interval of inferred R561H prevalence was created using a spatial Binomial logistic model.

Map figures were created in RStudio using spatial data downloaded from the DHS website.

## Results

By genotyping parasites from discarded DHS samples, we were able to investigate the distribution of diversity of *Pfk13* mutations in Rwanda early in the emergence of R561H.

### Short amplicon PCR was an effective way to determine infections

Amplification identified 25.4% of *Pf* unknown samples positive (476/1873), similar to the expected 25.7% positivity found by DHS by RDT. On a per cluster level, we found that amplicon success had moderate correlation with the expected *P. falciparum* prevalence (Pearson coefficient = 0.357). However, some clusters fell above or below the expected value based on DHS data due to inherent variance from testing different subsets of individuals such as differences in average age (RDT was 18 years compared to 22 years for PCR). While PCR is normally more sensitive than RDT, sample degradation likely decreased our sensitivity and our assays did not leverage multicopy genes. Additionally, RDTs can be false positives due to the persistence of HRP2 protein despite clearance of the parasite.

### Targeted deep sequencing of amplicon revealed R561H and other *Pfkelch13* mutations

Of the 476 PCR positive samples, 351 (73.7%) were successfully sequenced, revealing a majority of wild type parasites (341/351, 97.03%) (Supplementary Table 1). One sample (0.27%) had a synonymous mutation (G533G) and 9 samples (2.70%) had nonsynonymous K13 mutations, including R561H (4), V555A (3), C532W (1), and G533A (1) (Supplementary Table 1).

The four R561H isolates were spatially clustered across three DHS clusters in the southeast part of the country, representing 5.47% of samples in Kirehe and 2.74% in Ngoma. (Figure 2a, Supplementary Figure 1, Supplementary Table 2). The Moran’s I value of spatial correlation of clusters with R561H was 0.133 (p=0.046), indicating significant spatial clustering. Additionally, a 97.5% confidence interval of inferred R561H prevalence based on the number of mutant isolates, sampling scheme, and geographic proximity to other isolates highlighted localization in the southeast, higher confidence of low or no prevalence in the southwest, but lower of confidence in unsampled northern regions (Figure 2b).

**Figure 2:**
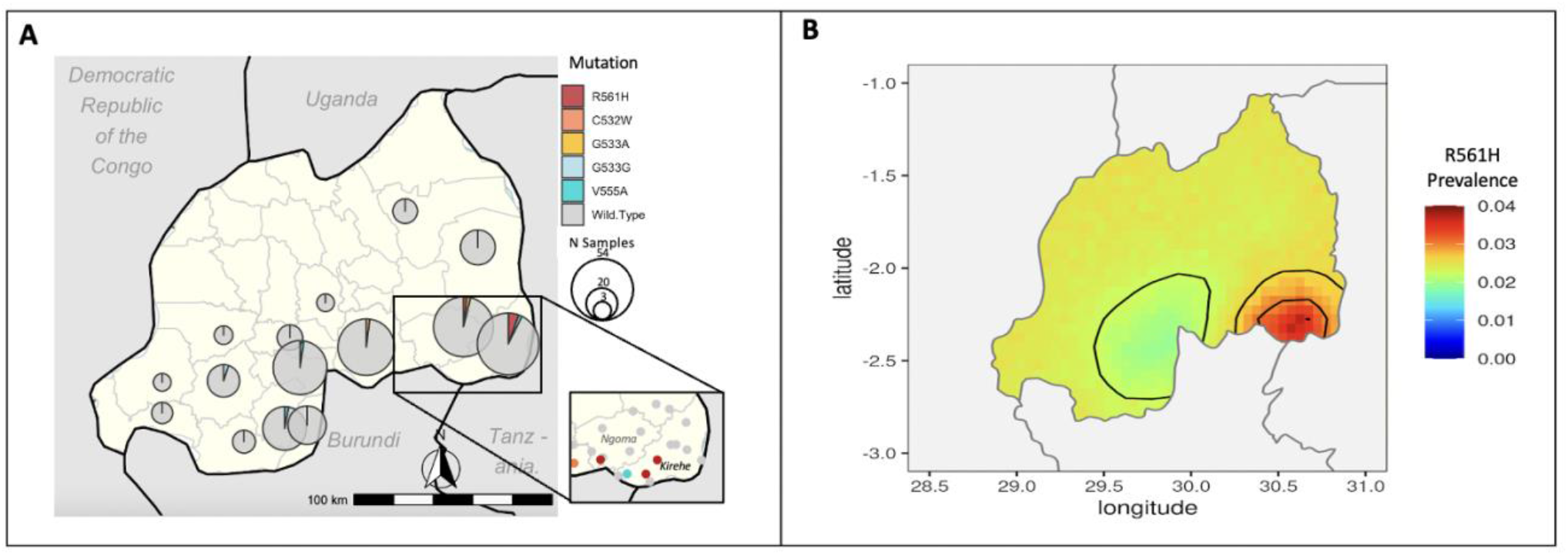
Geographic distribution of *Pfk13* mutations in high malaria prevalence demographic health survey clusters in Rwanda. **A:** Mutations confirmed by sequencing amplicons corresponding to *Pfk13* amino acids 516-572 are presented as pie charts, the center of each corresponding to the center of the DHS survey district. Mutation presence at the DHS cluster level shown for Kirehe and Ngoma districts. **B:** The 97.5% confidence interval of inferred R561H prevalence using a spatial Binomial logistic model.

## Discussion

Previous *Pfk13* genotyping in Rwanda in the years leading up to the first observation of R561H failed to capture the larger picture across the country. This study provides expanded geographic sampling and an improved baseline of *Pfk13* genotypes in relatively high-prevalence regions early in the R561H emergence timeline. Previous studies only observed the mutation in Masaka as of 2014, but our study demonstrates that it was also present and significantly clustered in high-transmission regions in the southeast of the country. In keeping with prior reports, we found a small number of clinically unvalidated polymorphisms, three previously not observed in Rwanda as of 2014-15 (Supplementary Table 3).

The presence of R561H in three spatially clustered high-transmission clusters outside of previously reported areas could point towards a more southern origin or spread. While additional data is needed to better understand the origin and spread, it is concerning that the resistant parasites were already present in high-transmission areas as of 2014. This was unexpected, as it is hypothesized that low-transmission areas drive selection of resistant parasites, due to low levels of host immunity and reduced within-host competition [15]. A possible explanation of the current findings could be periods of low- and high-transmission due to intermittent indoor residual spraying in these regions. Regardless of cause, the presence of mutant parasites in high-transmission zones is of significant concern for the control of malaria.

Though these samples allowed for retrospective genotyping, working with old, discarded samples poses challenges. Some bags were not fully sealed, some had water damage, and the desiccant was exhausted in most sample bags. The DNA was thus variably degraded, forcing us to focus on just a short amplicon containing the R561H mutation site. While this did not capture all mutations of interest in Rwanda (i.e. C469F, P574L, and A675V), these could be pursued with additional amplicons [1,7,10].

Relatively low sequencing success can likely be attributed to selection of PCR step 1 positive amplicons, as any sign of a band was classified as positive. The presence of non-specific bands could have led to misidentification of positives at this step.

The sampling scheme should also be taken into consideration. This method led to samples from older patients, with sequenced samples from patients averaging 22 years. As factors such as pre-existing immunity are thought to have an impact on the evolution of parasites, the relative proportion of mutant parasites may be different from studies with a lower median age such as Uwimana et al 2020, who only took samples from children 1-14 years [1].

This approach requires significant manual labor for large studies. Sorting through thousands of samples, punching dried blood spots, and running gels to identify *Plasmodium* positive samples was not a small task. This in practice required us to sample from high malaria prevalence areas to not waste resources and time on unknown samples in low-prevalence areas for little output. This is a key limitation as low-prevalence are historically the emergence sites of resistant parasites, so it is possible that R561H existed in 2014-15 in areas we did not sample. Finally, we did not sample directly from Masaka, where R561H was first observed in 2014. However, our sampling scheme included multiple clusters nearby Masaka and did not find R561H. When possible, future national sampling should coordinate HIV and RDT sampling for ease of genotyping residual samples.

Our study is the first country-wide examination of early distribution of artemisinin resistant *Pfk13* R561H parasites in Rwanda using samples derived from a representative sample of circulating parasites. This data will allow for better modeling of the initial rate of spread when combined with other retrospective analyses and current surveys.

## Footnotes

The authors declare no conflicting interests.

This laboratory work and sequencing using residual dried blood spot samples from the Rwanda Demographic and Health Survey 2014-15 was supported by NIH R01AI156267 to JAB.

Sequence files are accessible under the accession number PRJNA885045.

## References

1. Uwimana A, Legrand E, Stokes BH, et al. Emergence and clonal expansion of in vitro artemisinin-resistant Plasmodium falciparum kelch13 R561H mutant parasites in Rwanda. Nat Med. 2020; 26(10):1602–1608.

2. Asua V, Conrad MD, Aydemir O, et al. Changing Prevalence of Potential Mediators of Aminoquinoline, Antifolate, and Artemisinin Resistance Across Uganda. J Infect Dis. 2021; 223(6):985–994.

3. Straimer J, Gandhi P, Renner KC, Schmitt EK. High Prevalence of Plasmodium falciparum K13 Mutations in Rwanda Is Associated With Slow Parasite Clearance After Treatment With Artemether-Lumefantrine. J Infect Dis. 2022; 225(8):1411–1414.

4. Ashley EA, Dhorda M, Fairhurst RM, et al. Spread of Artemisinin Resistance in Plasmodium falciparum Malaria. N Engl J Med. 2014; 371(5):411–423.

5. Ariey F, Witkowski B, Amaratunga C, et al. A molecular marker of artemisinin-resistant Plasmodium falciparum malaria. Nature. 2014; 505(7481):50–55.

6. Takala-Harrison S, Jacob CG, Arze C, et al. Independent Emergence of Artemisinin Resistance Mutations Among Plasmodium falciparum in Southeast Asia. J Infect Dis. 2015; 211(5):670–679.

7. Bergmann C, Loon W van, Habarugira F, et al. Increase in Kelch 13 Polymorphisms in Plasmodium falciparum, Southern Rwanda. Emerg Infect Dis. 2021; 27(1):294–296.

8. Loon W van, Oliveira R, Bergmann C, et al. In Vitro Confirmation of Artemisinin Resistance in Plasmodium falciparum from Patient Isolates, Southern Rwanda, 2019. Emerg Infect Dis. 2022; 28(4):852–855.

9. Uwimana A, Umulisa N, Venkatesan M, et al. Association of Plasmodium falciparum kelch13 R561H genotypes with delayed parasite clearance in Rwanda: an open-label, single-arm, multicentre, therapeutic efficacy study. Lancet Infect Dis. 2021; 21(8):1120–1128.

10. Tacoli C, Gai PP, Bayingana C, et al. Artemisinin Resistance–Associated K13 Polymorphisms of Plasmodium falciparum in Southern Rwanda, 2010–2015. Am J Trop Med Hyg. 2016; 95(5):1090–1093.

11. National Institute of Statistics of Rwanda, Rwanda, DHS Program, editors. Rwanda demographic and health survey, 2014-15: final report. Kigali, Rwanda: Rockville, Maryland, USA: National Institute of Statistics of Rwanda, Ministry of Finance and Economic Planning: Ministry of Health ; The DHS Program, ICF International; 2016.

12. Makunin A, Korlević P, Park N, et al. A targeted amplicon sequencing panel to simultaneously identify mosquito species and Plasmodium presence across the entire Anopheles genus. Mol Ecol Resour. 2022; 22(1):28–44.

13. Aydemir O, Janko M, Hathaway NJ, et al. Drug-Resistance and Population Structure of Plasmodium falciparum Across the Democratic Republic of Congo Using High-Throughput Molecular Inversion Probes. J Infect Dis. 2018; 218(6):946–955.

14. Hathaway NJ, Parobek CM, Juliano JJ, Bailey JA. SeekDeep: single-base resolution de novo clustering for amplicon deep sequencing. Nucleic Acids Res. 2018; 46(4):e21–e21.

15. White NJ. Antimalarial drug resistance. J Clin Invest. 2004; 113(8):1084–1092.

