## Supplementary Table 1 for "Examining the early distribution of the artemisinin-resistant *Plasmodium falciparum* kelch13 R561H mutation in Rwanda"

**Supplementary Table 1: *Pfk13* haplotypes observed in sequenced samples (N=351)**

| **Mutation** | **Amino Acid Sequence** | **N Samples** | **Percent of Population (Weighted)** |
| --- | --- | --- | --- |
| Wild Type | 516-DVWYVSSNLNIPRRNNCGVTSNGRIYCIGGYDGSSIIPNVEAYDHRMKAWVEVAPLN-572 | 341 | 97.03% |
| C532W | 516-DVWYVSSNLNIPRRNNWGVTSNGRIYCIGGYDGSSIIPNVEAYDHRMKAWVEVAPLN-572 | 1 | 0.28% |
| G533A | 516-DVWYVSSNLNIPRRNNCAVTSNGRIYCIGGYDGSSIIPNVEAYDHRMKAWVEVAPLN-572 | 1 | 0.30% |
| G533G | 516-DVWYVSSNLNIPRRNNCGVTSNGRIYCIGGYDGSSIIPNVEAYDHRMKAWVEVAPLN-572 | 1 | 0.27% |
| V555A | 516-DVWYVSSNLNIPRRNNCGVTSNGRIYCIGGYDGSSIIPNAEAYDHRMKAWVEVAPLN-572 | 3 | 0.78% |
| R561H | 516-DVWYVSSNLNIPRRNNCGVTSNGRIYCIGGYDGSSIIPNVEAYDHHMKAWVEVAPLN-572 | 4 | 1.34% |
