## Supplementary Figure 1 for "Examining the early distribution of the artemisinin-resistant *Plasmodium falciparum* kelch13 R561H mutation in Rwanda"

### Slide 1
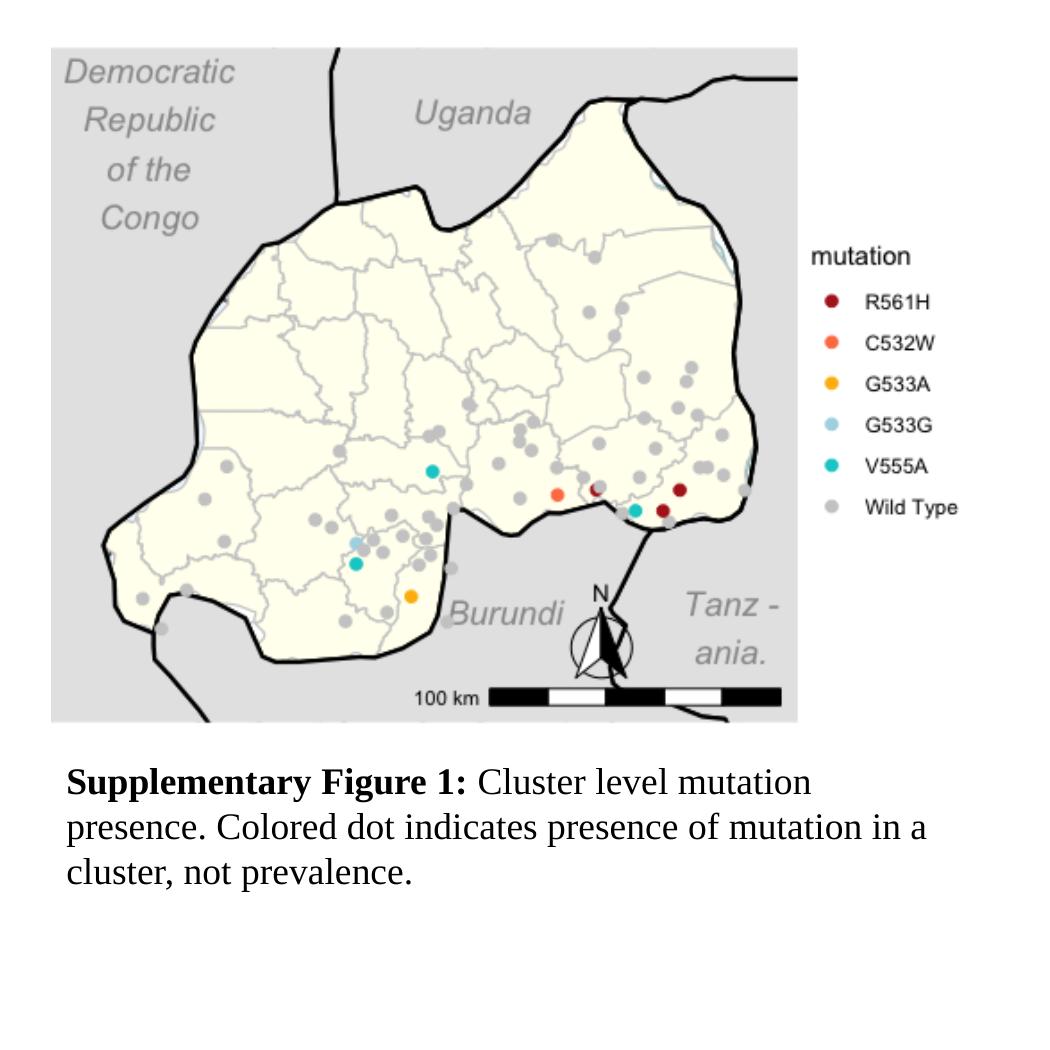

Supplementary Figure 1: Cluster level mutation presence. Colored dot indicates presence of mutation in a cluster, not prevalence.
