## Supplementary Methods for "Examining the early distribution of the artemisinin-resistant *Plasmodium falciparum* kelch13 R561H mutation in Rwanda"

Rebecca Kirby^1*^, David Giesbrecht^1*^, Corine Karema^2,3^, Oliver Watson^4,5^, Savannah Lewis^1^, Tharcisse Munyaneza^6^, Jean De Dieu Butera^6^, Jonathan J Juliano^7*^, Jeffrey Bailey^1*^, Jean-Baptiste Mazarati^8*^

^1^Brown University, Providence, RI, USA ^2^Quality Equity Health Care, Kigali, Rwanda, ^3^Swiss Tropical and Public Health Institute, University of Basel, Switzerland, ^4^MRC Centre for Global Infectious Disease Analysis, Imperial College London, London, UK, ^5^London School of Hygiene and Tropical Medicine, ^6^National Reference Laboratory, Rwanda Biomedical Center, ^7^UNC, Chapel Hill, NC, ^8^INES-Ruhengeri, Ruhengeri, Rwanda

**PCR Protocol**

**PCR step 1**

- Primers were manufactured by IDT and hydrated to 100 uM stock solutions in Low EDTA TE (Thermo #AAJ75793AP), then diluted to 10 uM in molecular grade water (MGW).
- DNA was centrifuged at 4000 rpm for 3 minutes before 2.5uL was transferred from the top of the tube to the reaction mixture to reduce chelex carryover.
- Non-template negative control and 3D7 positive controls were added to each plate.

**PCR step 2**

- 5uL of PCR product from PCR1 diluted 1:10 with molecular grade water were added immediately after dilution.

**Overall**

- Thermocycling was run on an Eppendorft Mastercycler 96
- PCR step 1 primers contained linkers and PCR step 2 primers contained unique barcodes to allow for multiplexing.

**PCR Step 1 Master Mix**

| **Reagents**  **(NEB #M0491, Thermo**  **BP28191)** | **Stock Concentration** | **Volume 1x reaction** |
| --- | --- | --- |
| **Molecular Grade Water** | NA | 11.75 uL |
| **Q5 Reaction Buffer** | 5x | 5 uL |
| **dNTPs** | 10 mM | 0.5 uL |
| **Primer F** | 10 uM | 2 uL |
| **Primer R** | 10 uM | 0.5 uL |
| **Q5 Polymerase** | 2000 U/mL | 0.25 uL |
| **MgCl** | 50 mM | 3 uL |
| **Total** |  | **22.5 uL** |
|  |  | +2.5 uL DNA |

**Primer Design**

| Forward Primer | 5’-GACTCGCCAAGCTGAAGNNNNCATAGCTGATGATCTAGGGG-3’ |
| --- | --- |
| Reverse Primer | 5’-ACGTGTGCTCTTCCGATCTNNNNCTGAGGTGTATGATCGTTTAAG-3’ |

**PCR Step 1 Thermocycling Parameters**

| **Cycle Step** | | **Temperature (C)** | **Time** | **No. Cycles** |
| --- | --- | --- | --- | --- |
| Initial Denaturation | | 98 | 30 seconds | 1x |
| Amplification | Denaturation | 95 | 10 seconds | 30x |
|  | Annealing | 56 | 15 seconds |  |
|  | Elongation | 72 | 20 seconds |  |
| Final Elongation | | 72 | 2 minutes | 1x |
| Hold | | 4 | Hold | 1x |

**PCR Step 2 Master Mix**

| **Reagents**  **(NEB #M0491, Thermo**  **#BP28191)** | **Stock Concentration** | **Volume 1x reaction** |
| --- | --- | --- |
| **Molecular Grade Water** | NA | 11.75 uL |
| **Q5 Reaction Buffer** | 5x | 5 uL |
| **dNTPs** | 10 mM | 0.5 uL |
| **Barcoded Primer F** | 10 mM | 1.25 uL |
| **Barcoded Primer R** | 10 mM | 1.25 uL |
| **Q5 Polymerase** | 2000 U/mL | 0.25 uL |
| **Total** |  | **20 uL** |
|  |  | +5 uL PCR step 1 product diluted 1:10 with MGW |

**PCR Step 2 Thermocycling Parameters**

| **Cycle Step** | | **Temperature (C)** | **Time** | **No. Cycles** |
| --- | --- | --- | --- | --- |
| Initial Denaturation | | 98 | 30 seconds | 1x |
| Amplification | Denaturation | 95 | 10 seconds | 6x |
|  | Annealing | 59 | 15 seconds |  |
|  | Elongation | 72 | 20 seconds |  |
| Final Elongation | | 72 | 2 minutes | 1x |
| Hold | | 4 | Hold | 1x |
