## Supplementary Table 2 for "Examining the early distribution of the artemisinin-resistant *Plasmodium falciparum* kelch13 R561H mutation in Rwanda"

**Supplementary Table 2: Survey weighted Pfk13 Mutation Prevalence Within Administrative Districts**

| **District** | **C532W** | **G533A** | **G533G** | **V555A** | **R561H** | **Wild Type** | **N Samples** |
| --- | --- | --- | --- | --- | --- | --- | --- |
| **Bugesera** | 1 (2.14%) | 0 | 0 | 0 | 0 | 47 | 48 |
| **Gatsibo** | 0 | 0 | 0 | 0 | 0 | 11 | 11 |
| **Gisagara** | 0 | 0 | 0 | 0 | 0 | 27 | 27 |
| **Huye** | 0 | 0 | 0 | 1 (3.01%) | 0 | 32 | 33 |
| **Kamonyi** | 0 | 0 | 0 | 0 | 0 | 3 | 3 |
| **Karongi** | 0 | 0 | 0 | 0 | 0 | 4 | 4 |
| **Kayonza** | 0 | 0 | 0 | 0 | 0 | 23 | 23 |
| **Kirehe** | 0 | 0 | 0 | 1 (1.78%) | 3 (5.47%) | 50 | 54 |
| **Ngoma** | 0 | 1 (1.63%) | 0 | 0 | 1 (2.74%) | 50 | 52 |
| **Nyamagabe** | 0 | 0 | 1 (5.37%) | 0 | 0 | 19 | 20 |
| **Nyamasheke** | 0 | 0 | 0 | 0 | 0 | 3 | 3 |
| **Nyanza** | 0 | 0 | 0 | 1 (1.93%) | 0 | 44 | 45 |
| **Nyaruguru** | 0 | 0 | 0 | 0 | 0 | 9 | 9 |
| **Ruhango** | 0 | 0 | 0 | 0 | 0 | 12 | 12 |
| **Rusizi** | 0 | 0 | 0 | 0 | 0 | 7 | 7 |
| **Total** | 1 | 1 | 1 | 3 | 4 | 341 | 351 |
