## Supplementary Table 3 for "Examining the early distribution of the artemisinin-resistant *Plasmodium falciparum* kelch13 R561H mutation in Rwanda"

**Supplementary Table 3: additional Pfk13 mutation information**

| **Mutation** | **In vitro artemisinin resistance status** | **Date First Observed, Location (Percent), Study** | **Other Observations in Rwanda: Date, Location (Percent), Study** | **Notes** |
| --- | --- | --- | --- | --- |
| C532W | Unvalidated | 2014-15, Bugesera (2.14%), **this study** | none |  |
| G533A | Unvalidated | 2014-15, Ngoma (1.63%), **this study** | - 2019, Huye District (1.5%), Bergmann et al 2021 | Associated with treatment failure in India, but cannot be directly linked to ACT resistance (Mishra et al 2015) |
| G533G | Unvalidated | 2014-15, Nyamagabe (5.37%), **this study** | none |  |
| R561H | Validated | 2014-15, Masaka (7.4%), Uwimana et al 2020 | - 2014-15, Kirehe (5.47%), Ngoma (2.47%), **this study** - 2018, Masaka (19.6%), Rukara (22%), Uwimana et al 2021 - 2019, Huye District (4.5%), Bergmann et al 2021 |  |
| V555A | Unvalidated | 2012, Huye District (1.2%), Tacoli et al 2015 | - 2014-15, Huye (3.01%), Kirehe (1.78%), Nyanza (1.93%), **this study** - 2015, Masaka (0.39%), Rukara (1.2%), Uwimana et al 2020 - 2019 Huye District (1.5%), Bergmann et al 2021 |  |
